# Accurate calling of low-frequency somatic mutations by sample-specific modeling of error rates

**DOI:** 10.1101/2024.12.17.629019

**Authors:** Yixin Lin, Carmen Oroperv, Peter Sørud Porsgård, Amanda Frydendahl Boll Johansen, Mads Heilskov Rasmussen, Thomas Bataillon, Mikkel Heide Schierup, Claus Lindbjerg Andersen, Kristian Almstrup, Søren Besenbacher

## Abstract

Calling rare somatic variants from NGS data is more challenging than calling inherited variants, especially if the somatic variant is only present in a small fraction of the cells in the sequenced biopsy. In this case, having a good estimate of the error rate of a specific base in a particular read becomes essential. In paired-end sequencing, where some DNA fragments are shorter than twice the read length, the overlapping regions of the read pairs are an ideal resource for training models to discern context-dependent base error rates, as any discordant bases in the overlaps must be caused by a sequencing error or an alignment error. We have created a new tool named BBQ (an acronym for Better Base Quality) that uses overlapping reads to estimate the error rate conditional on the mutation type, sequence context, and base quality. We also estimate how much the error rate of concordant bases in overlapping reads is decreased compared to bases in non-overlapping reads. Results show that overlapping reads can remove sequencing errors induced by DNA damage and that the increased quality of overlapping reads differs between samples and mutation types, reflecting different damage patterns between samples. We use the error models to call rare somatic variants. Sequencing data from a testis biopsy and a cell-free DNA sample serve as a proof-of-concept for rare germ cell mutation calling and for detecting rare cancer mutations. We find that using the sample-specific error models of BBQ allows us to call rare somatic variants with fewer false positives than existing tools such as Mutect2 and Strelka2.

## 1. Introduction

The genomes in the cells of an individual are not completely identical - postzygotic mutations start accumulating in cells after fertilization, and the genetic difference between cells gradually increases with age. Most research into postzygotic mutations has focused on the positively selected driver mutations involved in cancer development, but the identification of somatic mutations is becoming increasingly relevant in understanding the pathogenesis of non-cancerous diseases as well (Olafsson and Anderson 2021; Cagan et al. 2022; Martincorena and Campbell 2015). The vast majority of non-cancerous postzygotic mutations are only present in a small number of cells, and even in a very localized tissue biopsy, the mutation will thus only be present in a few sequence reads (Dou et al. 2018). This rarity of the mutations makes them hard to detect.

Most somatic variant callers focus on finding variants with variant allele frequencies higher than 5% (Cibulskis et al. 2013; Ma et al. 2019; Z. Chen et al. 2020), which is sufficient for finding most driver mutations in tumor biopsies but generally inadequate for detecting non-cancerous postzygotic mutations as well as possible future drug resistance mutations. Focusing on variants with frequencies over 5% is also insufficient for liquid biopsy analyses where the circulating tumor DNA (ctDNA) in blood is heavily diluted by the much larger amount of cell-free DNA (cfDNA) originating from healthy cells.

Comprehensive and accurate molecular profiles called from sequential cfDNA samples can enhance cancer research through sensitive liquid biopsy applications such as monitoring the tumor burden(Zviran et al. 2020; Widman et al. 2024), early recurrence prediction(Gale et al. 2022) and understanding the mechanisms of acquired drug resistance(Lheureux et al. 2023). These analyses, especially in a low tumor burden setting where the allele frequency in the cfDNA is low, rely on accurate variant detection to appropriately guide downstream clinical decisions. Correspondingly, accurate low-frequency variant calling could also improve the understanding of postzygotic non-cancerous diseases. According to the MosaicBase database, which is curated from publications from 1998-2018, there are over 250 non-cancer diseases related to postzygotic single-base mutations(Yang et al. 2020). Accurate variant detection from localized tissue samples is essential for the diagnosis of these diseases as well as the discovery of new disease-related variants.

The two primary sources of erroneous variant calls from NGS data are alignment and sequencing errors. Many existing variant calling methods focus on reducing alignment errors by realigning reads during variant calling and assembling reads into haplotypes (Cibulskis et al. 2013; Cooke, Wedge, and Lunter 2021; Kim et al. 2018). For rare variants, sequencing errors are, however, often a more significant problem.

The sequencing errors observed from Illumina machines - the predominantly used sequencing platform - can have different causes, including PCR errors during bridge amplification, crosstalk between adjacent clusters, and phasing problems between cycles (Ledergerber and Dessimoz 2011). Some nucleotides and sequence contexts are more prone to such errors than others (Stoler and Nekrutenko 2021; Nakamura et al. 2011; Frydendahl, Rasmussen, et al. 2024). The presence of DNA lesions in the input fragment can drastically increase the error rate during bridge amplification as some nucleotide modifications, such as 5-methylcytosine (5mC) and 8-oxo-guanine (8oG), can cause the polymerase to insert a wrong base (Arbeithuber, Makova, and Tiemann-Boege 2016). Oxidative damage to DNA during library preparation can, in particular, lead to a high number of errors when an 8oG erroneously pairs with an adenine, resulting in a C→A error (Costello et al. 2013).

One strategy to reduce the impact of sequencing errors is to use sequence data from a set of normal (non-cancerous) samples to learn which positions or sequence contexts are most error-prone. Shearwater (Gerstung, Papaemmanuil, and Campbell 2014) uses a set of independent training samples to train a position-specific error model based on the observed rate of read alignment. DREAMS-vc (Christensen et al. 2023) uses training samples to build a read-level error model in order to calculate the likelihood that a putative mutation is caused by sequencing errors. For these methods that rely on training samples to work optimally, it is necessary to have a large number of training samples collected, processed, and sequenced using exactly the same approach. This is problematic when there are technical or biological differences between the available samples or when only one sample is available.

Another strategy to reduce sequencing errors is to use specialized library preparation approaches that tag each initial molecule, thus allowing the formation of high-quality consensus sequences. Duplex sequencing (Schmitt et al. 2012), which tags each strand of the initial molecule independently, is capable of producing ultrasensitive NGS data, as the independent consensus sequences for the forward and reverse strands enable the removal of errors coming from DNA lesions, present in only one strand as well as PCR errors created during cluster formation. However, Duplex sequencing and later improvements to the method, such as NanoSeq (Abascal et al. 2021), are costly, as a large number of reads are needed to obtain the final consensus sequence of one initial DNA molecule (Arbeithuber, Makova, and Tiemann-Boege 2016). These methods also require large amounts of starting material, limiting their use on cfDNA and other clinical samples with limited material.

To address the challenge of calling rare mutations from standard NGS data without a matching set of training samples, we present a novel bioinformatical method called BBQ (short for Better Base Qualities), specifically designed to improve the detection accuracy of rare somatic mutations from NGS data. The method uses overlapping reads to train sample-specific sequencing error rate models and then uses these error rate models to call rare somatic variants. We describe the development and validation of BBQ by presenting its key features and comparing its performance to established tools on both a cfDNA and a non-cancerous somatic mutation data set. The results demonstrate that BBQ achieves superior precision in identifying rare somatic mutations.

## 2. Results

### 2.1 Discordant bases in overlapping reads

In paired-end sequencing of DNA fragments shorter than twice the read length, there will be an overlapping portion of the read-pair (Figure 1). The main idea of BBQ is that discordances between the overlapping reads must be the consequence of an error (either a sequencing error or an alignment error). This means that such discordant base pairs can be used as good-quality training data for a model that estimates the probability that a specific base in a read is wrong. Figure 1 shows the steps implemented in BBQ and explained in turn below and in detail in the Methods section.

**Figure 1.**
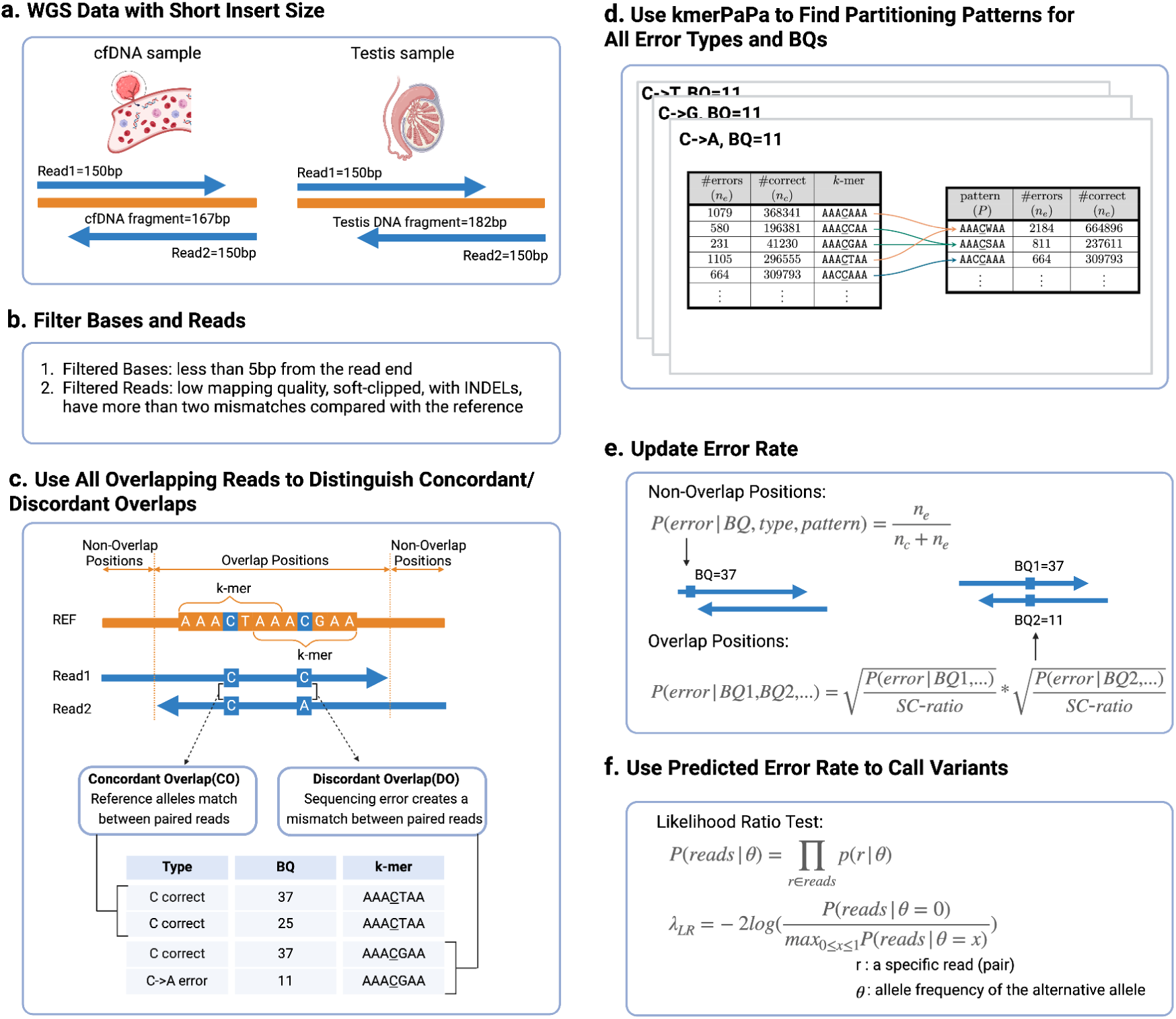
Overview of the workflow for BBQ variant calling. The variant calling process of BBQ consists of six main steps. **a.** Input data (cfDNA and testis data) consists of paired-end NGS data with a short average insert size. **b**. Reads that are more likely to contain errors are filtered as well as bases near the end of a read. **c.** At all positions where the individual is deemed to be homozygous for the reference allele, all overlapping positions are classified as a concordant overlap (CO) if the bases of the paired reads match or as a discordant overlap (DO) if they do not. Only overlaps containing at least one reference allele are used as training data. The alternative alleles in the discordant overlaps are assumed to be the result of an error, and each one is assigned to one of the 6 different strand-symmetric error types depending on the reference and alternative alleles. Bases matching the reference allele are assumed to be correct. Each base is additionally annotated with its base quality and the reference k-mer surrounding the position. **d.** For each combination of base quality and error type, kmerPaPa is used to find a set of k-mer patterns for which the error rates differ. **e.** For positions not in overlapping pairs, we calculate the probability of a specific type of error based on the pattern and the base quality. For overlap positions, we take the quality of both of the two bases into account and update the rate using an SC-ratio (Single alternative to Concordant overlapping alternatives ratio) (Supp. Figure 1) that reflects how often a specific error type is seen at non-overlap positions compared to concordant overlaps. **f.** Variant calling is performed using a likelihood ratio test (*λ_LR_*) that compares the likelihood of a variant if the true variant allele frequency, θ, is zero (and all alternative alleles must thus be due to errors) to the maximum likelihood if θ is allowed to be non-zero.

### 2.2 Low-frequency somatic mutation benchmark datasets

To evaluate BBQ’s accuracy, we used two benchmark data sets (Figure 1a). One is based on synthetically creating low-frequency somatic variants by reducing the allele frequency of inherited variants. The other is a cfDNA data set from a cancer patient, where sequencing data from a tumor biopsy is used to identify the true tumor mutations present in the cfDNA. The following two subsections explain the creation of these two benchmark data sets.

#### 2.2.1 Synthetic benchmark data from a testicular biopsy

The synthetic benchmarking data was created from a human testis biopsy sequenced with a mean depth of 140x and a mean insert size of 182 base pairs. The same sample was also sequenced with PacBio HiFi technology with a mean depth of 60x. Because the same biopsy was used for sequencing, the two data sets were expected to have high concordance for heterozygous variants. Therefore, we called heterozygous SNVs from the PacBio HiFi data and subsequently lowered the allele frequency of these variants in the Illumina data. Each alternative allele at the variant position was changed to the reference allele with a probability of 0.975, which resulted in a set of low-frequency variants with an average allele frequency of 0.018 (see methods section for details).

#### 2.2.2 Cell-free DNA benchmark data

The cfDNA benchmarking data was derived from Illumina whole-genome sequencing of DNA isolated from the plasma of a patient with relapsed colorectal cancer. The composite testing sample was generated by merging four post-operation (post-OP) cfDNA samples obtained from the same patient. This testing data exhibited a mean depth of 98X and a mean tumor fraction of 2.01%. To remove non-cancer somatic variants, a buffy coat sample from the same patient was sequenced to a mean depth of 41X. Non-reference variants seen in the buffy coat sample were filtered out from the somatic variant list of the cfDNA.

For validation, a matched tumor biopsy was collected from the same colorectal cancer patient and sequenced. We used Mutect2 to call somatic SNVs from the tumor data under tumor-normal mode, and the identified tumor SNVs were considered the true variants that we aimed to detect in the cfDNA dataset.

### 2.3 Filtering bases and reads

Somatic variant callers usually aim to deliver the best trade-off between false positives and false negatives. In this study, we focus on calling low-frequency mutations close to the detection limit. Recognizing that we will not be able to call all such low-frequency mutations, our main priority is to reduce the number of false positives. This focus on reducing false positives, even at the cost of some false negatives, is essential when we are calling variants close to the limit of detection, as the false positives could easily outnumber the true positives. In order to limit the number of false positives, BBQ uses strict filters to ignore alternative alleles in reads that might be aligned incorrectly or have low quality. We ignore all bases that are less than five base pairs from the end of a read since they are known to be more error-prone (Ma et al. 2019), and we disregard reads that have low mapping quality, are soft-clipped, contain indels, or have more than two mismatches with the reference (Figure 1b). In addition, a set of genomic regions that contain repeat sequences is disregarded from analysis to avoid variant calling at low-complexity regions.

### 2.4 Estimating error rates using sequence context

We use discordant bases in overlapping reads to learn which sequence contexts are more likely to produce sequencing errors of a specific kind. If we only look at the sites where we believe the proband to be homozygous for the reference allele, we can assume that the alternative allele in a discordant pair is the error and assign the error to one of six different types (A→C, A→G, A→T, C→A, C→G, C→T). At such sites where the proband is homozygous for the reference allele, we can similarly assume that all reference bases in overlapping reads are correct. Both the error bases and correct bases are annotated with their base quality and the reference 7-mer centered around the base (See Figure 1c). For each combination of error type and base quality, BBQ then applies the kmerPaPa algorithm (Bethune, Kleppe, and Besenbacher 2022) to partition the k-mers into patterns, thus avoiding overfitting the error rate of specific k-mers due to sparsity (Figure 1d). Figure 2a shows the predicted error rates for each pattern, base quality (BQ) and mutation type. These error rates reflect the probability of seeing a specific error in one read (not both members of an overlapping pair), and we thus use these error rates for bases outside overlapping regions.

**Figure 2.**
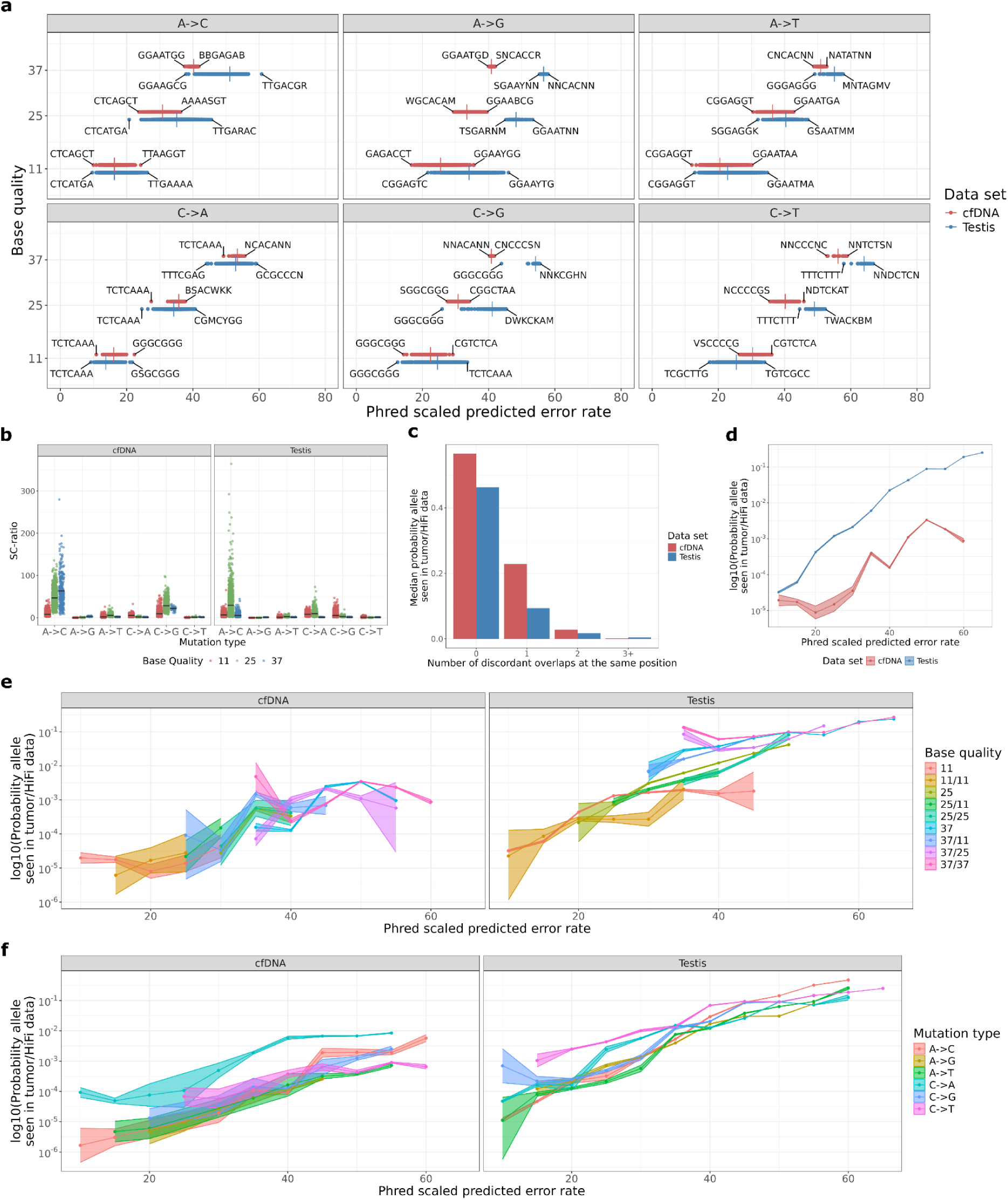
**a.** Phred scaled predicted error rates of 7-mer patterns. The patterns with the highest and lowest predicted error rates are indicated with text labels. Vertical lines indicate the mean error rates. **b.** The ratio between the rate of alternative alleles seen in non-overlapping and overlapping positions (SC-ratio). The data points indicate the SC-ratios of 7-mer patterns. Black horizontal lines show the mean SC-ratios. **c.** The median fraction of alternative alleles seen in the validation set for different numbers of discordant overlaps observed at the same position. The median fraction was calculated based on genomic positions where three or more alternative alleles were observed. **d, e, f** Fraction of alternative alleles seen in the validation set given the predicted error rate.

In overlapping regions, we check whether the alleles are concordant and otherwise ignore the reads when calling variants at this position. This means we expect a lower error rate for alleles in overlapping regions since the exclusion of apparent errors observed in discordant overlaps will remove some of the sequencing errors. To estimate how much lower the error rate is in concordant overlaps compared to non-overlapping regions, we counted the number of alternative alleles in both concordant overlaps and outside overlaps. We then calculate the ratio between the rate of alternative alleles seen in single non-overlapping and concordant overlapping bases to estimate how much lower we expect the error rate to be in overlapping reads for a specific mutation type, base quality, and k-mer pattern. We refer to this “Single alternative to Concordant overlapping alternatives ratio” as the SC-ratio (Supp. Figure 1), and the calculated values of this ratio can be seen in Figure 2b. As expected, most ratios are above one, indicating that we see more alternative alleles in non-overlapping positions compared to concordant overlapping positions. Furthermore, the results show significant differences between the datasets (Figure 2b). The cfDNA data set primarily shows elevated error rates at non-overlapping positions for A→C and C→G mutations. The effect is generally smaller in the testis data, and the mutation types with the largest ratios are A→C and C→A.

These differences between the error rates in overlapping and non-overlapping regions are likely due to differences in the type of DNA damage. Damage to the DNA is usually restricted to one strand and can result in an error if the sequenced strand contains the damage. In the overlapping reads where both strands are sequenced, the error will, however, create a discordant overlap, allowing us to remove the error. The elevated error rates for A→C and C→A mutations in non-overlapping regions of reads can be explained by oxidative damage. An oxidized guanine base, known as 8-oxo-guanine (hereafter 8OG), can result in a mismatched pairing with adenine, causing 8OG:A instead of 8OG:C and hence a C→A error. If an 8OG is misincorporated instead of T during replication, the result will be an A→C error (Costello et al. 2013; L. Chen et al. 2017). The differences in damage patterns between the two test samples might reflect the different natures of the data sets. The cfDNA data was fragmented *in vivo* by DNases and amplified using PCR before sequencing. In contrast, the testis data was fragmented in the lab using sonication followed by size selection using beads but did not go through PCR amplification. These differences in error patterns emphasize the importance of creating sample-specific error models trained on the test data itself.

### 2.5 Modeling position-specific error rates

Besides sequence context, several other factors that we are not taking into account can affect the likelihood of observing a specific type of sequencing error at a particular site. Looking at a site where an alternative allele is observed three times or more (not counting discordant overlaps), we observe that this alternative allele is less likely to be in the validation set if the site also contains a discordant overlap (Figure 2c). To reflect this, we use the error rate predicted based on k-mer, mutation type, and base quality as the prior rate and update it based on the number of discordant overlaps observed at the position in question using a Beta-Binomial distribution (see Methods).

### 2.6 Estimating validity of the predicted error rates

Figure 2d compares the Phred-scaled predicted error rate of an observed alternative allele to the probability that the allele is in the validation set. As expected, the fraction of alternative alleles seen in the validation set increases as the Phred-scaled error rate increases. The overall fraction of true alternative alleles is lower in the cfDNA data set, which can be explained by larger amounts of DNA damage and PCR-induced errors.

The error rates are predicted independently for each base quality, but we expect the higher base qualities to have lower error rates. Figure 2e compares the predicted error rates with the fraction of true alternative alleles separately for each possible base quality at non-overlap positions and each combination of base qualities at concordant overlap positions. Each base quality or combination of base qualities shows a range of predicted error rates and follows the expectation that alternative alleles with higher Phred-scaled predicted error rates are more likely to be true (a high value on the Phred scale corresponds to a low error probability). As expected, the lowest error rates and the highest fractions of true alternative alleles are observed for base qualities 37, 37/37 (overlap with two concordant BQ 37 alleles), and 37/25. It also shows that the predicted error rates are an improvement to the existing base qualities. We observe that some BQ 25 alleles are predicted to have error rates similar to BQ 37 alleles, whereas others have error rates more akin to BQ 11 variants, and the validation data show that those with the low error rates are much less likely to be errors than those predicted to have high error rates. Figure 2f compares the predicted error rates of the six mutation types. The error rates across all mutation types show a linear trend where alleles with higher Phred-scaled error rates are more likely to be observed in the validation set.

### 2.7 Calling variants using the error models

We use the estimated error rates to calculate the likelihood of observing a given set of reads at a position given the variant allele frequency at that position. This allows us to compare the maximum likelihood estimate of a variant to the null hypothesis that the variant allele frequency (VAF) of the variant is zero - i.e., the presence of the alternative allele is due to sequencing errors. We use this likelihood ratio to rank the possible variants in each test sample, employing a cutoff greater than 10, and compare the results to commonly used somatic variant callers (Figure 3). BBQ reaches higher precision values than the other methods in both the cfDNA and testis data sets. However, neither dataset can reach a recall close to 100% due to the low average variant allele frequency. We calculate the recall values based on the total number of variants in the validation set but some of the variants in the validation set are not seen at all in the alignment we call variants in. For the cfDNA data, the true alternative allele is not present for 39.5% of the variants, while it is absent for 18.8% of the variants in the testis data (Supp. Figure 2).

**Figure 3.**
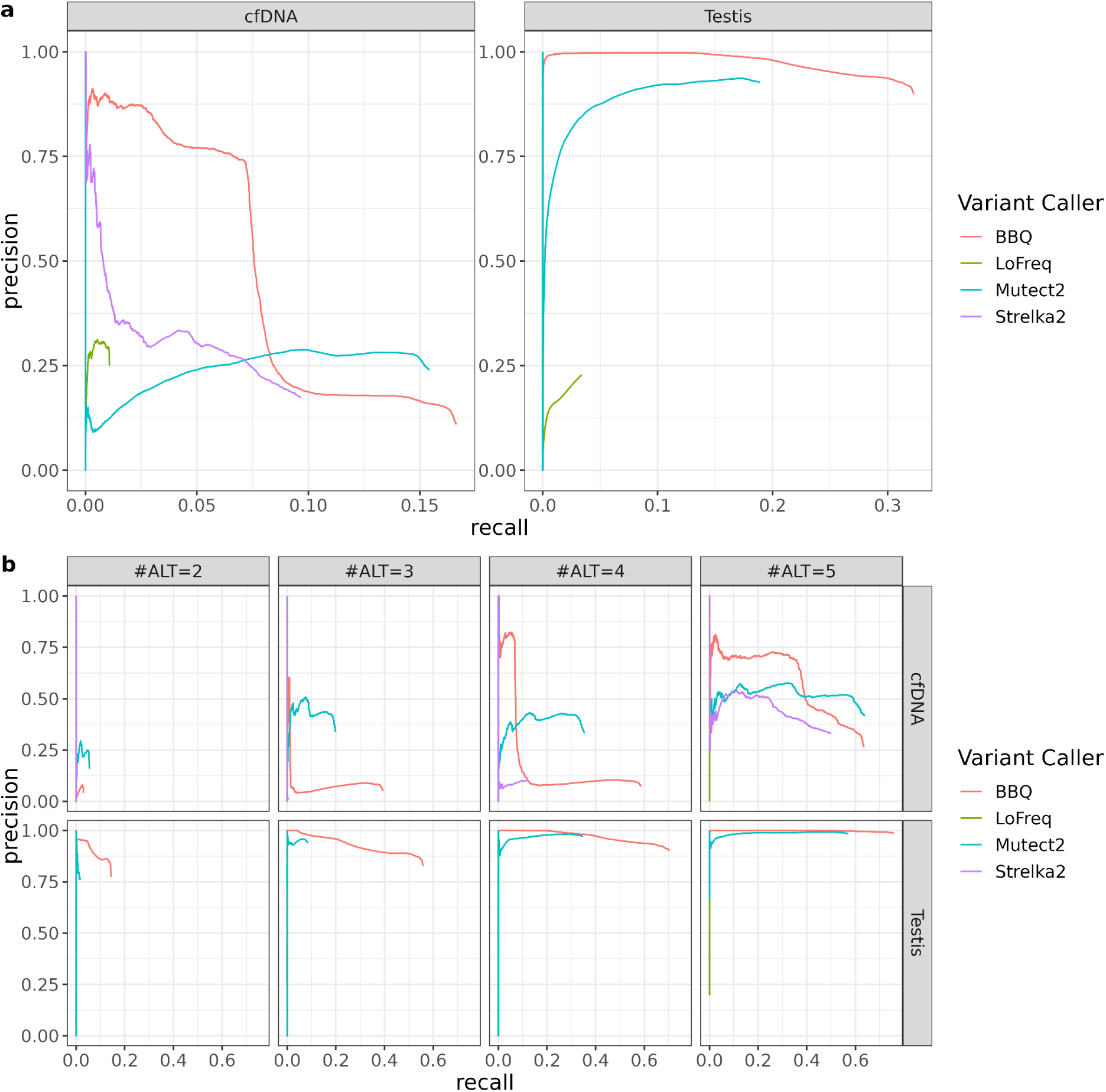
**a** Precision-Recall curves for both cfDNA and testis datasets. BBQ, LoFreq, Mutect2 and Strelka2 were used for cfDNA benchmarking, while BBQ, LoFreq and Mutect2 were used for testis data benchmarking. **b** Precision-Recall curves where the putative variants have been stratified based on the number of times the alternative allele is observed. #ALT=x refers to variants where the alternative allele is observed x times (without any filtering) in the bam file.

If we split the validation variant sets based on how many DNA fragments they are present in (at any base quality), we observe that detecting an allele twice is adequate to achieve a precision exceeding 75% in the testis dataset. Additionally, BBQ shows superiority in both recall and precision compared with Mutect2 (Cibulskis et al. 2013) and Lofreq (Wilm et al. 2012). The same trend is also observed for variants with alternative counts of 3, 4 and 5 in the testis dataset. In the cfDNA dataset, BBQ demonstrates an advantage in precision (up to 63%) for a subset of variants when the alternative allele is observed more than 3 times, compared to Strelka2 (Kim et al. 2018), Mutect2, and Lofreq, though with very limited recall. When the count of alternative alleles is increased to 5, BBQ exhibits higher precision (∼75%) than all other variant callers for 40% of recall.

The fact that we can call variants seen in just two DNA fragments in the testis data set, whereas we need to observe the alternative allele four times to reach a precision above 50% in the cfDNA data, likely reflects differences in the generation of the data sets. The presence of PCR errors as well as increased levels of DNA damage in the cfDNA data results in a higher level of noise in this data set compared to the testis data set.

Sample-specific differences in the error patterns are further highlighted when swapping the error models of the two data sets during variant calling (Supp. Figure 3). When the testis-specific error rate model is applied for variant calling in the cfDNA, the precision significantly decreases (Supp. Figure 3a). This suggests that some error patterns that are present in cfDNA are not accounted for in the testis-specific model but are crucial for accurate variant calling. On the contrary, the precision of variant calling from the testis data is not remarkably affected when applying the cfDNA error model, suggesting that the major error patterns of the testis data set that are relevant for variant calling are also represented in the cfDNA-specific model. Nevertheless, when the variant sets are split by the alternative allele count, both the cfDNA and testis datasets exhibit a loss in precision after swapping the error models (Supp. Figure 3b). Furthermore, there is a clear decrease in the recall rate of testis variants observed two to four times when applying the cfDNA model. These variations in error patterns across the two samples suggest the benefit of sample-specific error modeling and can be superior compared with modeling by a panel of normals (PON), especially for the cfDNA datasets prone to damage during library preparation.

## 3. Discussion

Our analyses demonstrate that BBQ can use discordant overlaps in read-pairs to create error models that accurately capture the patterns of errors in a specific sample. By using these sample-specific error models, BBQ can achieve higher precision for calling rare somatic variants than other somatic variant callers. The goal of BBQ is to call variants that are close to the limit of detection with Illumina data, and the focus is thus on being able to distinguish variants only present in a few reads from sequencing errors present in a few reads. Besides using predicted error rates based on the context and site of a putative variant, we also apply stricter filters on reads than many other variant callers to keep the false discovery rate low.

The newest Illumina sequencing machines do not use the full range of Phred scale values to represent base qualities; instead, they group base qualities into four different values. This grouping can reduce the ability to separate true alternative alleles from sequencing errors. We have demonstrated that the error rates calculated by BBQ can discriminate between good-quality and bad-quality alleles within the same base quality group. This ability to adjust base quality based on empirically proven errors in the sample is ultimately what is increasing the ability of BBQ to identify true alternative alleles.

A limitation of BBQ is that it requires a substantial number of read pairs containing overlapping segments to have enough data to reliably train a good error model. This means that if the average insert size is much larger than 2 times the read length, then there will not be enough overlapping segments to train good error models. Since the majority of cfDNA fragments have a length close to 167 bp (Jiang et al. 2015), there will always be a sufficient number of overlapping segments in such data to train error models. As the calling of rare cfDNA variants is crucial to detect minimal residual disease after surgery or to diagnose early-stage tumors, such data sets are an obvious use-case for BBQ. To make optimal use of BBQ on other data sets, it will, however, often be necessary to deliberately adapt the usual sequencing protocol for the data set to achieve shorter-than-usual insert sizes. Furthermore, from the comparison of the two data sets in this study, it seems advisable to avoid PCR amplification in the library preparation as this is likely a major error source in the cfDNA data set. However, in situations where it is only possible to have very low amounts of input DNA, this might not be possible.

Overall, BBQ demonstrates advantages over existing variant callers when it comes to calling somatic variants present in very low frequencies. This ability to call rare mutations with good precision is essential for reliably detecting mutations in healthy tissue, subclonal mutations in cancers, or ctDNA mutations in patients with early-stage cancers. BBQ puts a higher weight on specificity than sensitivity compared to most existing variant callers. BBQ will thus especially be useful in situations where the frequencies of mutations in a sample are so low that it will not be possible to detect all mutations in the sample, but BBQ will still be able to correctly call a subset of the mutations without also calling a lot of false positives. The detection of (subsets of) mutations with good specificity is relevant both in basic research studies investigating somatic mutation processes as well as in clinical research where e.g. detection of circulating tumor DNA in plasma can be used in the clinical diagnostic workup.

## 4. Methods

### 4.1 Test data sets

#### 4.1.1 Synthetic benchmark data

The synthetic benchmark data set was created from a human testis sample obtained from a postmortem autopsy of an anonymous donor who consented to donate tissue for research purposes. DNA was isolated using the MagAttract HMW DNA kit (Qiagen, Hilden, Germany) and sequenced on an Illumina NovaSeq with a mean depth of 140x and a mean insert size of 182 base pairs. The DNA from the same testis sample was also sequenced with PacBio HiFi technology with a mean depth of 60x on a Sequel II instrument using 8M SMRT cells (Pacific Biosciences). As the same biopsy was used for sequencing, the two data sets were expected to have high concordance for heterozygous variants. Therefore, we called heterozygous SNVs from the PacBio HiFi data and subsequently changed the allele frequency of these variants in the Illumina data (see below). The long-read (average 14302 bp) HiFi data was also used to create a *de novo* haploid genome assembly, used as a reference genome for read alignment. After adapter removal with cutadapt, the Illumina reads were aligned to the haploid assembly with BWA-MEM. This was followed by applying samtool’s *fixmate* command to fill in the read mate information and tag duplicate reads with samtool’s *markdup* command. Supplementary Figure 4 shows an overview of the data set generation workflow.

Variant calling from the aligned HiFi reads was carried out using the same strategy as in (Porsborg et al. 2024). Only assembly contigs longer than 10^6^ bases were considered for variant calling. At each reference position, reads that had a mapping quality equal to 60 were arranged in a pileup format, and the variant was called if a set of requirements was fulfilled. Briefly, the variant was called only if one alternative allele was observed across all aligned reads, sequencing depth was higher than the 5th percentile (19x) and lower than the 99.7th percentile (73x), both the reference and the alternative allele had an allele frequency above 10% to avoid spurious artifacts, less than 10% of the aligned reads contained an indel, the variant position was further than 15 bp from the closest indel position, the variant was not in a repeat region and the position was not surrounded by homopolymer sequences longer than six bases. Indel positions were defined as positions where nucleotide bases made up less than 90% of the sequencing depth. A variant was defined to be in a repeat region if the number of unique 4-mers extracted from the reference 32-mer surrounding the variant position was below 16. The length of the homopolymer sequences surrounding the position was defined as the sum of the lengths of the preceding and succeeding homopolymers. All reference genome positions where an SNV was not called due to the abovementioned requirements were saved and excluded from the analysis when evaluating variant calling results on the Illumina data.

Positions of the heterozygous SNVs called from HiFi data were modified in the Illumina data by changing the majority of the alternative alleles to reference alleles. This was done by arranging Illumina reads to a pileup format and iterating over reads that were aligned to the given SNV position. If an alternative allele was observed, it was changed to the reference allele with a probability of 0.975. If a read-pair with alternative alleles was overlapping at a given position, reads from the pair were considered for modification together to ensure that no additional discordant overlapping positions in read-pairs were created. These changes resulted in a set of low-frequency variants with an average allele frequency of 0.018. Variant calling from the data with lowered allele frequencies was also carried out with BCFtools and all variants called with allele frequencies above 10% were excluded from the analysis.

To compare variant calling performance, Mutect2, and LoFreq tools were used to call variants on the Illumina data. Default parameters were used for both tools. Mutect2 was run in tumor-only mode, followed by variant filtering with the FilterMutectCalls command.

#### 4.1.2 Cell-free DNA benchmark data

The cell-free DNA benchmarking data originates from a colorectal cancer patient who experienced clinical recurrence. Four post-operation (post-OP) cell-free DNA samples from the patient were whole-genome sequenced using NovaSeq6000. The tumor fractions in the four samples estimated by MRDetect (Zviran et al. 2020), were 6.84%, 0%, 0% and 0%, with corresponding mean depths of 29.5X, 27.6X, 20.3X and 23.1X, respectively. The benchmarking data was created by merging the whole-genome sequencing data of the four samples, creating a pooled sample with a mean depth of 98X and a mean tumor fraction of 2.01%.

DNA from a matched fresh frozen tumor biopsy, obtained from the same colorectal cancer patient, was also whole-genome sequenced on NovaSeq6000 with a mean depth of 40X. DNA isolated from a buffy coat sample collected from the same patient was also sequenced on NovaSeq6000, yielding a mean depth of 41X.

All FASTQ data was aligned to hg38 and marked with duplicates by samtools. We used Mutect2 to call somatic mutations from the tumor data with the matched buffy coat sample. The workflow included running Mutect2 in tumor-normal mode, followed by LearnReadOrientationModel, CalculateContamination, and FilterMutectCalls steps. The identified 42063 SNVs were regarded as the validation sets that we aimed to detect from the cell-free DNA dataset (See Supp. Figure 5a).

To compare variant calling performance, Mutect2, Strelka2 and LoFreq were selected to call variants from the cfDNA merged data (See Supp. Figure 5b). All the tools were run under the tumor-normal mode with default setups. Mutect2 calls were additionally filtered by the FilterMutectCalls command.

### 4.2 Filtering reads, bases and positions

BBQ uses a set of strict quality filters to reduce the number of false positive variant calls. Sequenced bases that are positioned less than five bases from the end of the read are ignored. In addition, sequencing reads that have a mapping quality below 50, are soft-clipped, contain indels or have more than two mismatches with the reference genome are also disregarded. Lastly, a set of low-complexity genomic regions are ignored during k-mer counting and variant calling to avoid spurious artifacts that may arise due to repeat sequences. All reference genome regions that were annotated as ‘Simple_repeat’, ‘Low_complexity’ or ‘Satellite’ by RepeatMasker (Smit, Hubley, and Green, n.d.) were ignored. For the testis dataset, we ran RepeatMasker version 4.1.5 on the *de novo* genome assembly. For the cfDNA dataset, we downloaded the hg38 RepeatMasker track from the UCSC Genome Browser.

### 4.3 Counting errors and correct bases

We only use sites where we believe the proband to be homozygous for the reference allele as training data. This way, when we see a discordant overlap consisting of an alternative base and a reference base, we can assume that the alternative base is the erroneous base. We then annotate each erroneous base with the reference k-mer centered at the position and its base quality (11, 25 or 37 in Novaseq data). If the reference allele is not either cytosine(C) or adenine(A) we use the reverse complement of the reference allele and k-mer so that we only have 6 different strand-symmetric error types (C→A, C→G, C→T, A→C,A→G, A→T) and 2 different correct base types (C, A). We use, *n_e_^x→Y,q,m^*, to denote the total number of *X→Y* errors within each k-mer context, *m*, and base quality, *q*. We assume that all reference bases in overlaps at the sites used for training are correct and denote the total number of correct *X* bases with a specific combination of base quality, *q*, and k-mer, *m*, as *n_c_^X,q,m^*.

### 4.4 Training k-mer models

We denote the probability that a read erroneously contains the base *Y* if the real base is *X* as *e_X→Y_*. We want to calculate this probability conditional on the reference k-mer around the position and the BQ of the erroneous base. If k (the length of the k-mer) is large, we can only expect a few observed discordant overlaps for a specific k-mer, resulting in a large uncertainty in the error estimate. To avoid this, we use a k-mer pattern partitioning algorithm, kmerPaPa (Bethune, Kleppe, and Besenbacher 2022), to group the k-mers into a set of IUPAC patterns and only estimate a distinct rate for each pattern. This method partitions the set of k-mers using a set of IUPAC patterns so that each k-mer will match one and only one pattern *P*, while making sure that k-mers with similar error rates are grouped together. The number of errors and correct bases for a specific pattern can then be calculated by summing over the k-mers matching the pattern.

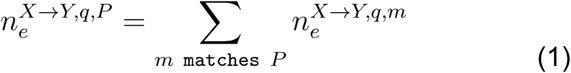

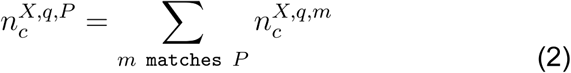

The estimated rate of X to Y errors for bases with quality q at sites that match the k-mer pattern, P, can then be calculated as:

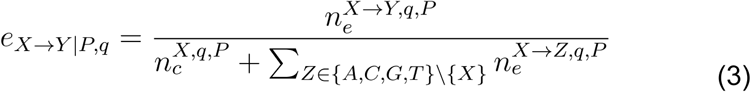

### 4.5 Handling overlapping reads

The formula (3) above gives an estimate of the probability of an error at bases not in the overlap between two reads from the same pair. We do not trust discordant bases in overlapping read pairs, and we ignore such bases during variant calling. For concordant bases in the overlap of a read pair, we want to calculate a joint error rate based on the two bases. If the two reads in a read pair were independent, we could calculate the joint error rate by multiplying the error rates of the two reads. Our tests, however, show that this would underestimate the error rate. Instead, we estimate a correcting factor by measuring how much more confident we are for bases in overlaps. We refer to this correction factor as the “Single alternative to Concordant overlapping alternatives ratio” abbreviated to “SC-ratio”, and we calculate it conditional on the mutation type, base quality, and k-mer pattern. The rate of single alternatives is calculated based on the observed number of *Y* alleles at *X* reference sites with base quality, *q*, that match k-mer pattern, *P*, at non-overlapping positions 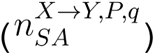 and the number of reference alleles at such sites 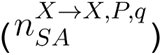:

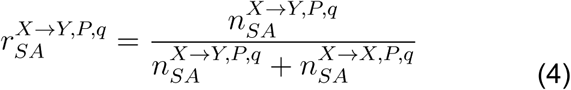

Similarly, we can calculate the rate of concordant *Y* alternative alleles at *X* reference sites where two paired reads overlap and both reads contain a *Y* allele with quality, *q*, using the number of concordant alternative variants 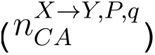 and the number of concordant reference variants 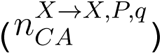:

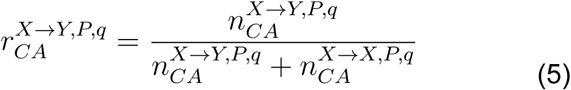

The SC-ratio (Supp. Figure 1) can then be calculated as the ratio between the rate of single alternative alleles and the rate of concordant alternative alleles for a specific base quality and k-mer pattern:

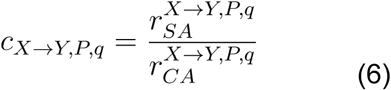

The error rate for concordant bases in an overlapping read pair where both bases have quality, *q*, is then calculated by dividing the estimated error rate for non-overlapping positions by the SC-ratio:

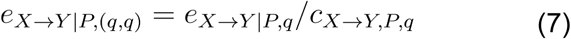

For concordant overlapping bases with different qualities, *q*_1_ and *q*_2_, we then calculate the joint error rate as the geometric mean of the error rate for (*q*_1,_ *q*_1_) pairs and (*q*_2_, *q*_2_) pairs:

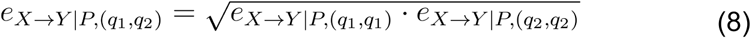

This corresponds to an arithmetic mean on the log (and Phred) scale and is preferable to the arithmetic mean of the probabilities where the resulting mean would be dominated by the highest error rate.

### 4.6 Updating error probabilities with position-specific information

We use Bayes’ formula to update the expected rate of *R* to *A* errors, *e_R→A_*, based on the number of discordant *R/A* overlaps, *x*, and the number of concordant *R/R* overlaps, *y*, at the site. Where R is the reference allele, and A is an alternative allele.

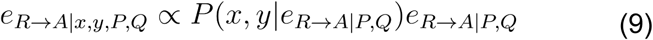

Q denotes the base quality of the alternative allele, either as a single value or as a tuple if the allele is in an overlapping read pair.

We assume that the prior probability of an error is beta-distributed

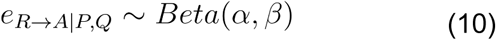

To parametrize the prior distribution, we choose a mean (mu) that matches the estimated error rate based on the base quality and k-mer combination, as described in sections 4.4 and 4.5. Besides the mean, we also introduce a hyperparameter, *N_prior_*. In our context, *N_prior_* can be interpreted as a pseudo sample size underlying the beta distribution, that modulates the weight we put on the prior distribution. From *μ* and *N_prior_*, we can calculate the shape parameters of the beta distribution as *α* = *μ* * *N_prior_* and β = *N_prior_* − *α*.

We used the binomial distribution to calculate the likelihood of x and y, given the prior error rate:

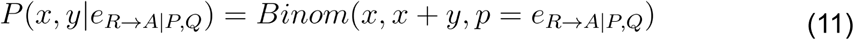

The posterior then follows an updated beta distribution:

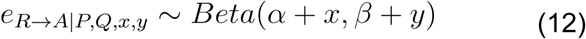

The weight of the prior, *N_prior_*, is a parameter to the variant calling command. Low values of *N_prior_* will give a low quality to alternative alleles at sites where discordant overlaps are observed, and this tends to increase the precision but also reduce the recall slightly. We have set the default value of *N_prior_* to 50000, which improves the precision (compared to not doing the positional update) while only having a slight effect on the recall (Supp. Figure 6).

The posterior error rate now differs between sites based on the number of discordant overlaps observed at the site, s. Since the reference k-mer is the same at a given site, we can also write the error probability conditional on the site, *s*, and base quality(s), *Q*: *e_R→A|s,Q_*.

### 4.7 Calculating the probability of not seeing an error

Section 4.3-4.6 explains how we calculate *e_X→Y|s,Q_* where *X* ≠ *Y*. To also calculate the probability of not seeing an error, *e_X→X|s,Q_*, we use the idea that the rates from a specific base should sum to 1, so we can calculate it as:

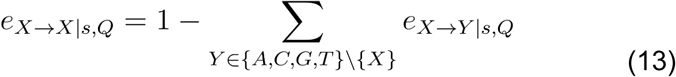

### 4.8 Calling variants using Likelihood Ratio Test

When we call a particular alternative allele, *A*, at a position, *s*, we only consider that alternative allele and the reference allele, *R*. Any read that contains a third allele is disregarded. The likelihood of a specific read (or read pair if the two reads overlap at position *s*), *r*, containing allele, *X_r_* ɛ {*A, R*} with quality *Q_r_* (a tuple or a single value depending on whether the two reads overlap at position *s*) at the position in question can then be written as a mixture distribution:

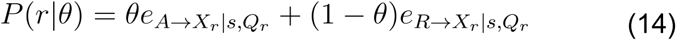

Where the unknown mixture proportion, *θ*, corresponds to the allele frequency of the alternative allele. The formula above assumes that the reads are mapped correctly. If we also consider the probability that the read is mapped to the wrong position, *p_map,r_* we can write the probability as:

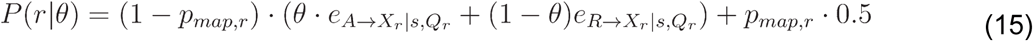

This assumes that if the read is erroneously mapped, then the probability that it carries the alternative allele is 50%, since it must carry either the alternative or reference allele to be considered.

We calculate the likelihood of all the reads containing either the reference allele or the alternative allele at position s, given the allele frequency, as:

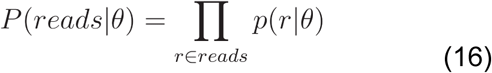

We can then compare the hypothesis that an alternative allele with an unknown frequency exists to the null hypothesis that no such allele exists using a likelihood ratio test statistic:

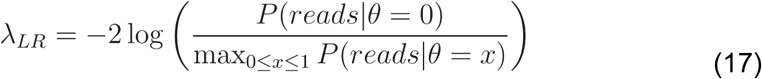

### 4.9 Filtering putative variants

As in Mutect, we filter putative variants where the median BQ of the alternative alleles is less than 20 by default. We also filter variants where the read filters remove significantly more alternative alleles than reference alleles based on Fisher’s exact test.

#### 4.9.1 Filtering based on sequencing depth

All variants with the filter “PASS” are further filtered based on the filtered depth and total depth distributions of these variants. The filtered depth at each position is calculated during variant calling after filtering out reads that have more than two mismatches to the reference, include indels or read clippings, have mapping quality below 50, or where the distance to the read end from the given position is shorter than five base pairs. The total depth is calculated as the total number of reads aligning to the given position without excluding reads based on the above-mentioned filters.

For depth-based filtering of variants, the 5% and 95% quantiles of the PASS variants’ filtered depth distribution and total depth distribution are calculated. Subsequently, BBQ filters all PASS variants with a filtered depth or total depth value below the respective 5% quantiles or above the respective 95% quantiles.

## Data availability

The sequenced testicular DNA originates from an anonymous donor and according to Danish legislation, we are not allowed to deposit human sequencing data from anonymous human donors. The sequence data from the analyzed cfDNA, tumor biopsy, and matching normal is from (Frydendahl, Nors, et al. 2024). Details about the data and how to apply for access can be found at: https://genome.au.dk/library/GDK000005/. Access to clinical data and processed sequencing data output files used in the article requires that the data requestor (legal entity) enter into Collaboration and Data Processing Agreements, with the Central Denmark Region (the legal entity controlling and responsible for the data). Request for access to raw sequencing data furthermore requires that the purpose of the data re-analysis is approved by The Danish National Committee on Health Research Ethics.

## Code availability

The BBQ source code is publicly available at https://github.com/BesenbacherLab/BBQ and a snakemake workflow to run the software is available at https://github.com/BesenbacherLab/bbq_pipeline.

## Acknowledgments

We acknowledge funding from the Novo Nordisk Foundation(grants number NNF21OC0069105 and NNF21OC0069056) and the Danish Cancer Society (grant number R257-A14700).

## Supplementary Figures

**Supp. Figure 1.**
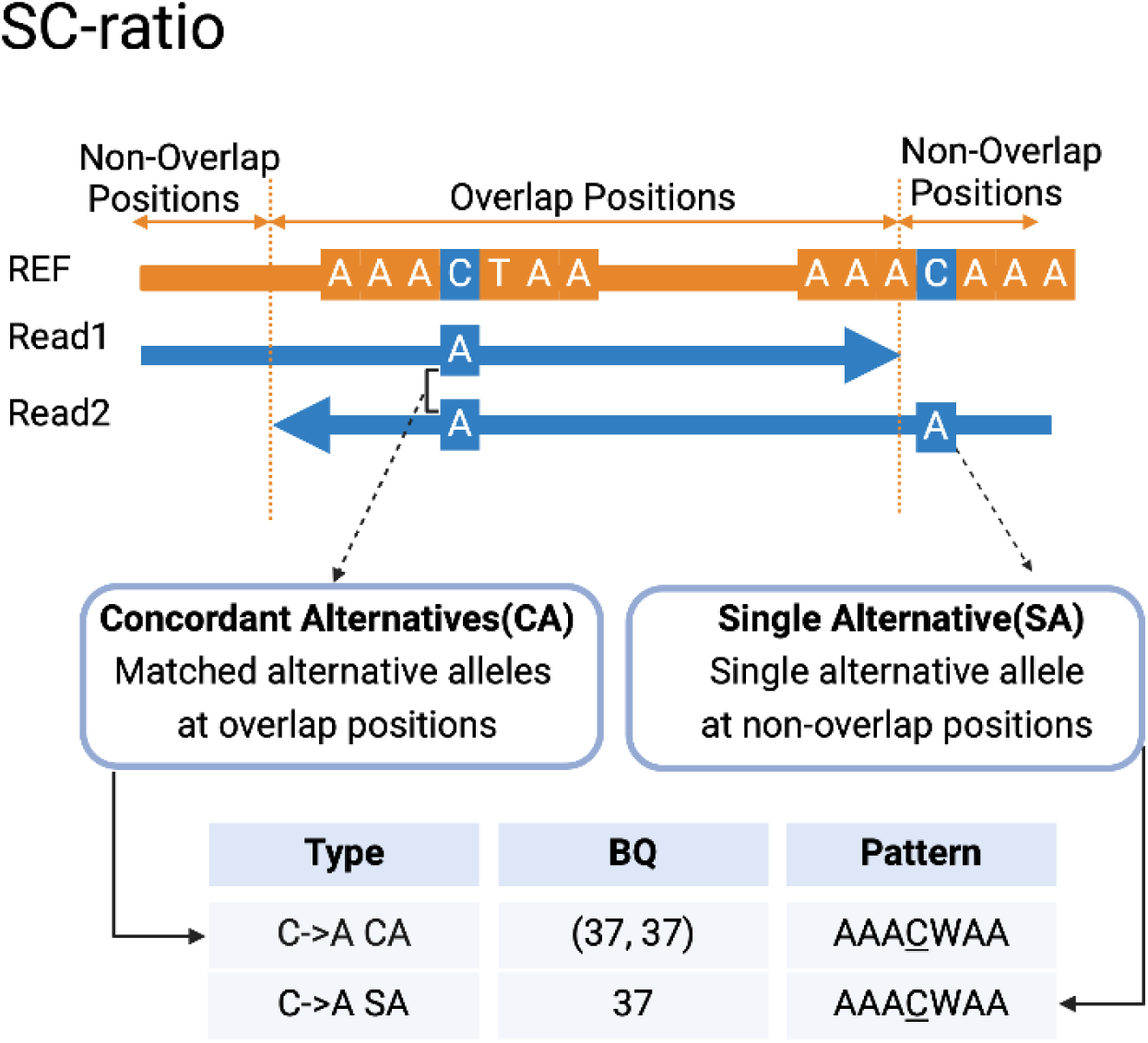
Illustration of the method used to distinguish concordant alternatives (CA) from single alternatives (SA) for calculating the SC-ratio. The counts of CA and SA are stratified by type, base quality (BQ), and pattern.

**Supp. Figure 2.**
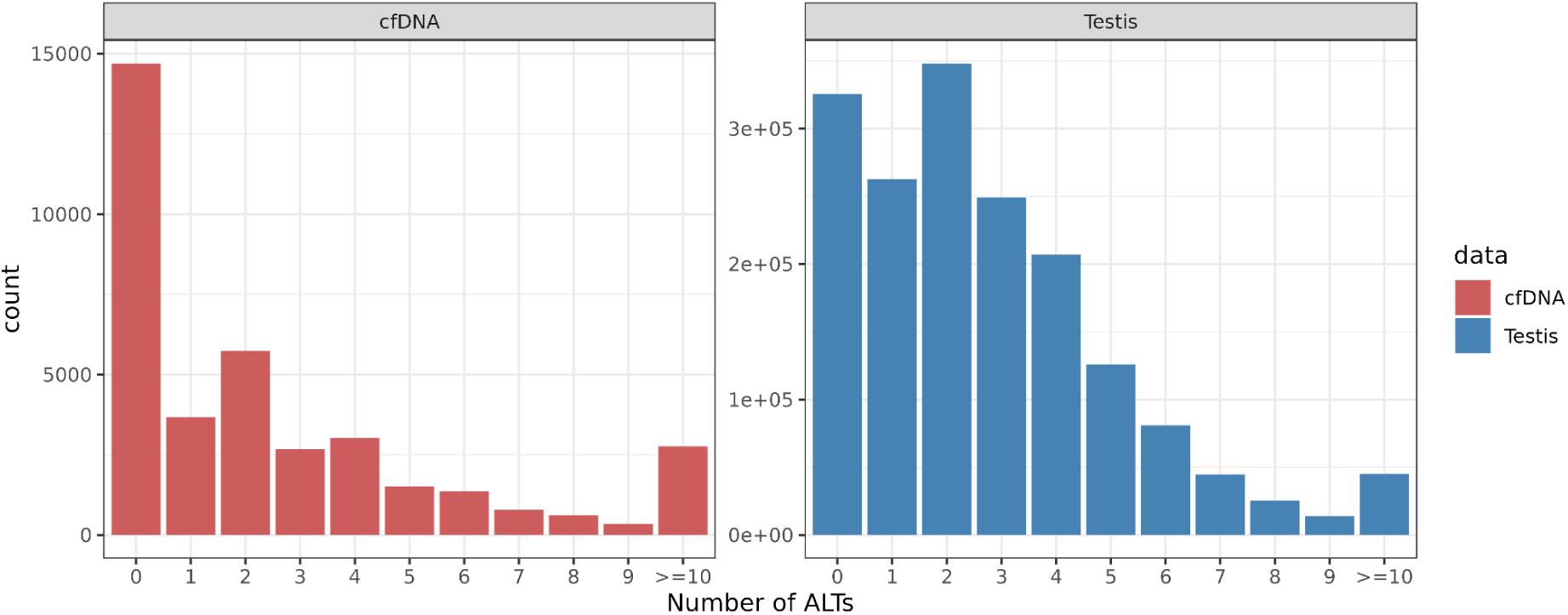
The distribution of the number of alternative alleles (ALTs) for the validation sets of cfDNA and testis data. In the cfDNA dataset, 14696 out of 37200 (39.5%) alternative alleles are not detectable. In the testis dataset, 325588 out of 1728868 (18.8%) alternative alleles are not present.

**Supp. Figure 3.**
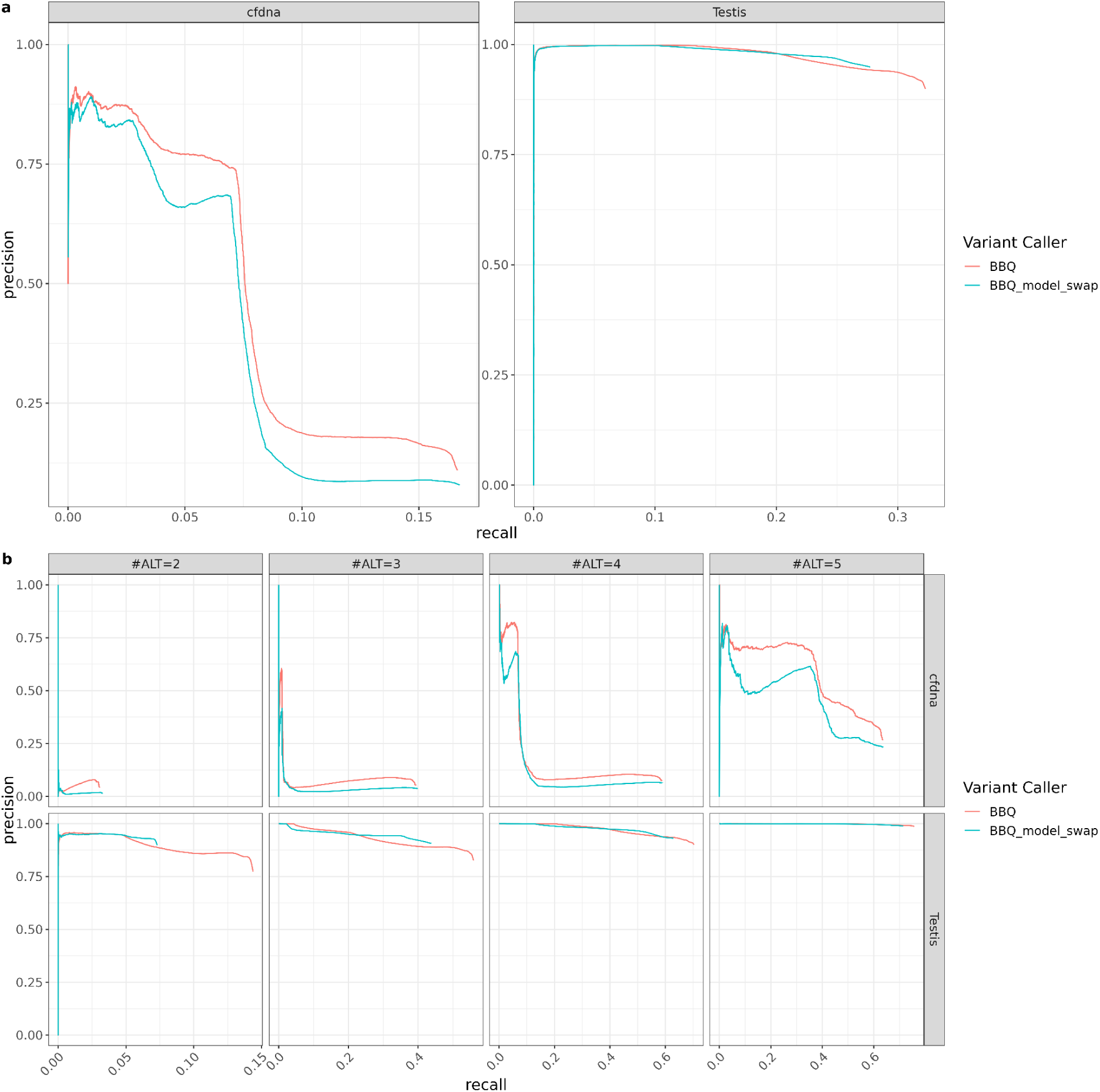
**a** Precision-Recall curves for both cfDNA and testis datasets. ‘BBQ’ refers to results obtained using the sample-specific model, whereas ‘BBQ_model_swap’ refers to results from the swapped model, where cfDNA data was analyzed using the testis model, and testis data were analyzed using the cfDNA model. **b** Precision-Recall curves stratified by specific alternative counts (2-5) for both the sample-specific BBQ model and sample-swapping BBQ model.

**Supp. Figure 4.**
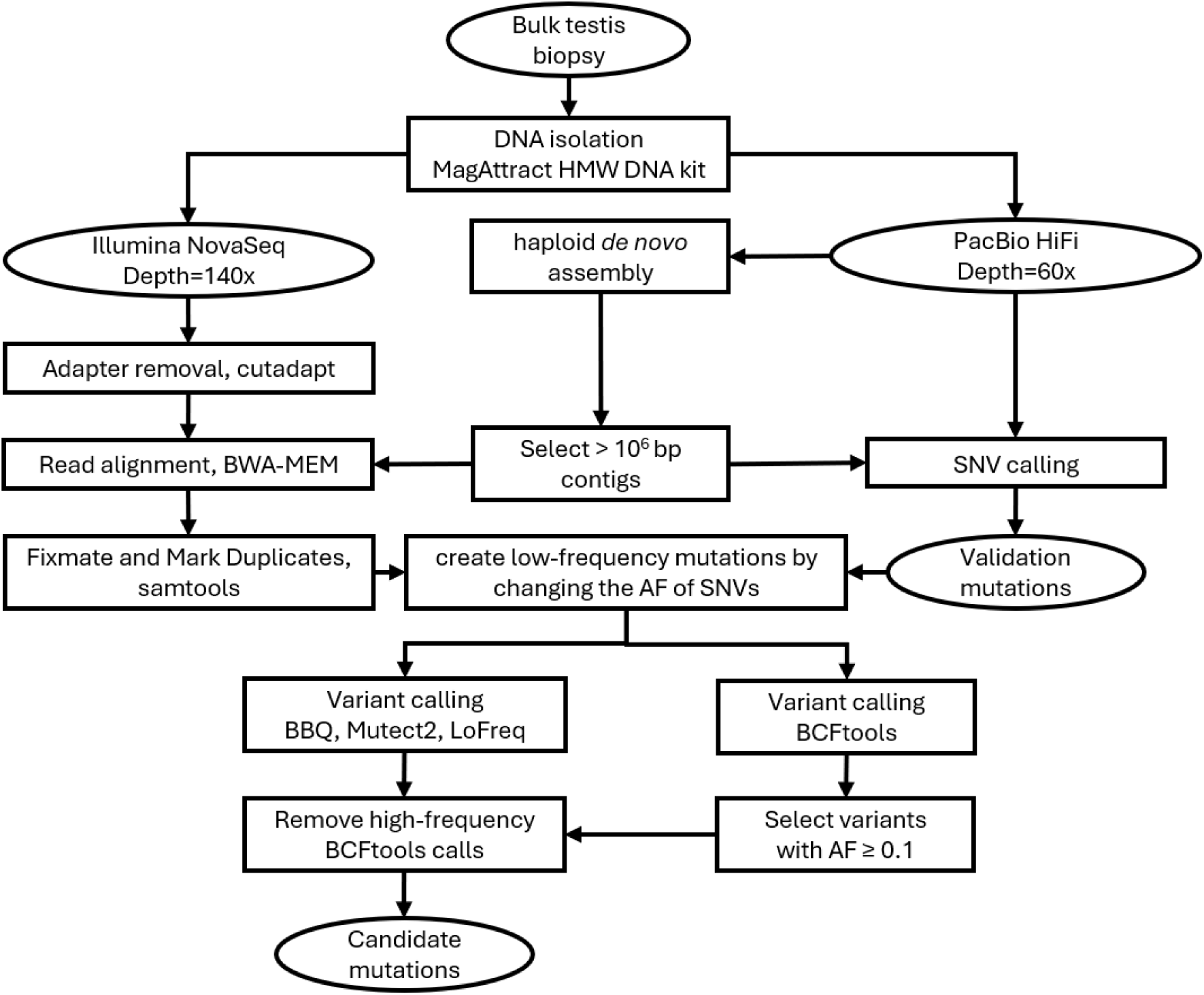
Procedures for generating synthetic benchmark data from a testis sample. The workflow illustrates the generation of the validation mutation set from PacBio HiFi data set and steps for somatic variant calling from Illumina data set. AF=allele frequency, bp= base pair.

**Supp. Figure 5.**
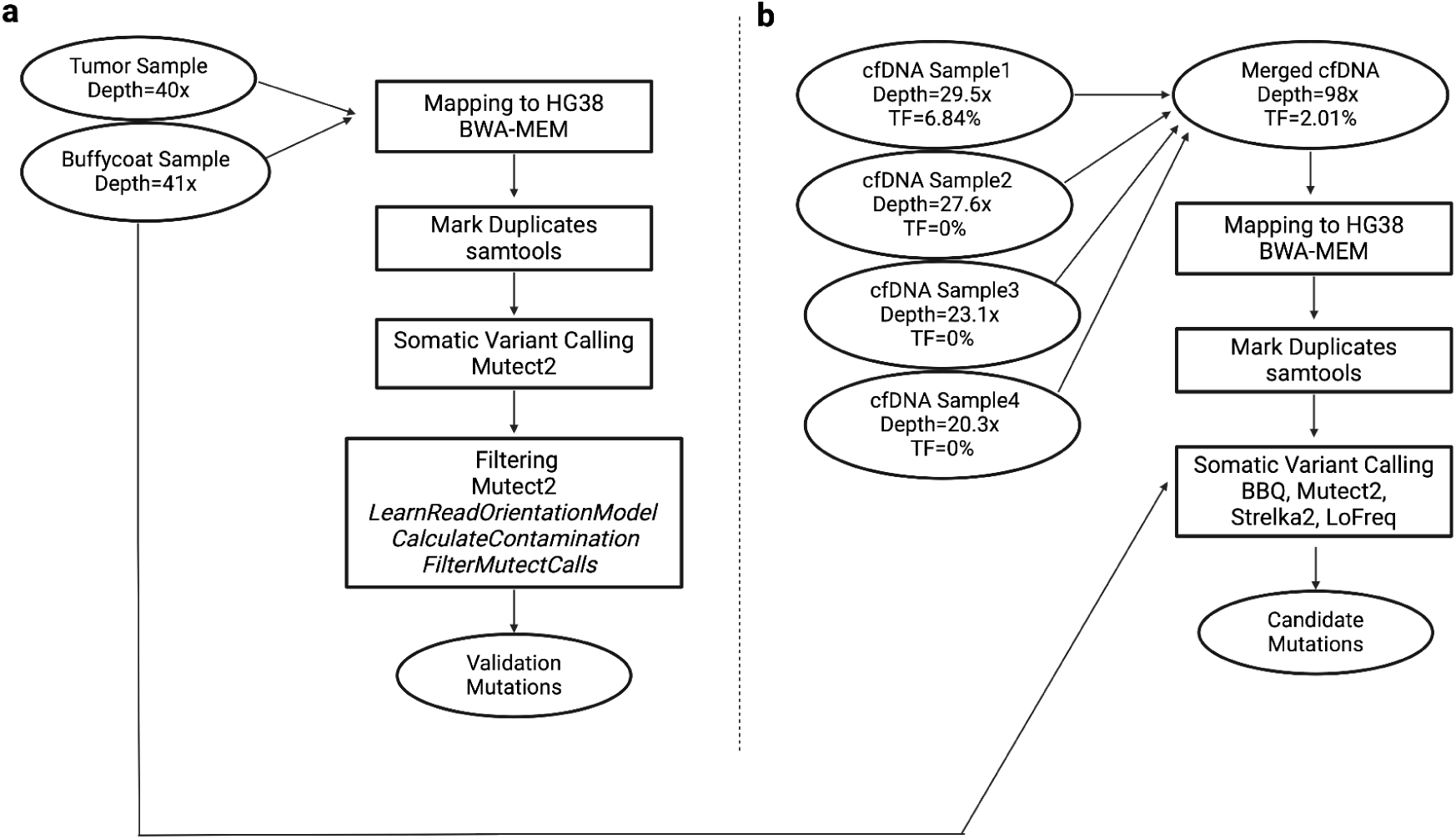
Procedures for generating cell-free DNA benchmarking data. **a.** Overview of the generation of validation mutation sets through somatic variant calling for paired tumor and buffy coat samples by Mutect2. **b.** Overview of the somatic variant calling process for the merged cell-free DNA sample. The tools BBQ, Mutect2, Strelka2, and LoFreq were selected for benchmarking.

**Supp. Figure 6.**
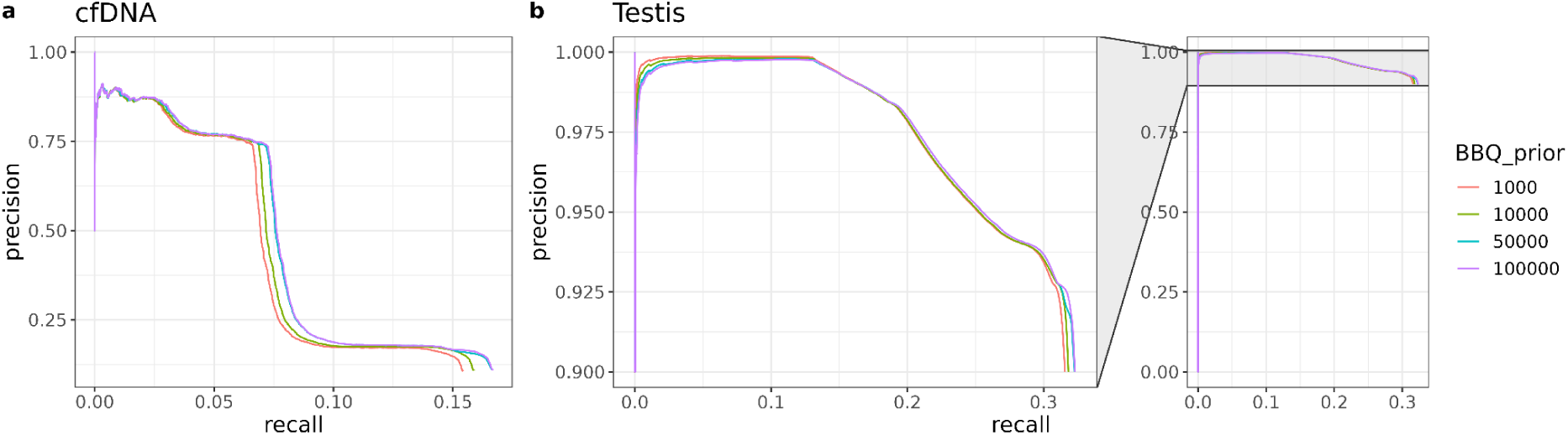
Comparison of different priors applied in a beta-binomial model to update error probabilities using position-specific information for variant calling.

